# Understanding the phase separation characteristics of nucleocapsid protein provides a new therapeutic opportunity against SARS-CoV-2

**DOI:** 10.1101/2020.10.09.332734

**Authors:** Dan Zhao, Weifan Xu, Xiaofan Zhang, Xiaoting Wang, Enming Yuan, Yuanpeng Xiong, Shenyang Wu, Shuya Li, Nian Wu, Tingzhong Tian, Xiaolong Feng, Hantao Shu, Peng Lang, Xiaokun Shen, Haitao Li, Pilong Li, Jianyang Zeng

**Affiliations:** Institute for Interdisciplinary Information Sciences, Tsinghua University, Beijing, China; Beijing Advanced Innovation Center for Structural Biology, Beijing Frontier Research Center for Biological Structure, Tsinghua University-Peking University Joint Center for Life Sciences, School of Life Sciences, Tsinghua University, Beijing, China; Silexon AI Technology Co., Ltd., Nanjing, Jiangsu Province, China; Bioinformatics Division, BNRIST/Department of Computer Science and Technology, Tsinghua University, Beijing, China; Protein Preparation and Identification Facility, Technology Center for Protein Science, Tsinghua University, Beijing, China; Institute of Pathology, Tongji Hospital, Tongji Medical College, Huazhong University of Science and Technology, Wuhan, Hubei Province, China; Convalife (Shanghai) Co., Ltd., Shanghai, China; Beijing Advanced Innovation Center for Structural Biology, Tsinghua-Peking Joint Center for Life Sciences, Department of Basic Medical Sciences, School of Medicine, Tsinghua University, Beijing, China; MOE Key Laboratory of Bioinformatics, Tsinghua University, Beijing, China

## Abstract

The ongoing coronavirus disease 2019 (COVID-19) pandemic has raised an urgent need to develop effective therapeutics against the severe acute respiratory syndrome coronavirus 2 (SARS-CoV-2). As a potential antiviral drug target, the nucleocapsid (N) protein of SARS-CoV-2 functions as a viral RNA chaperone and plays vital and multifunctional roles during the life cycle of coronavirus^1-3^. In this study, we discovered that the N protein of SARS-CoV-2 undergoes liquid-liquid phase separation (LLPS) both *in vitro* and *in vivo*, which is further modulated by viral RNA. In addition, we found that, the core component of the RNA-dependent RNA polymerase (RdRp) of SARS-CoV-2, nsp12, preferentially partitions into the N protein condensates. Moreover, we revealed that, two small molecules, i.e., CVL218 and PJ34, can be used to intervene the N protein driven phase separation and loosen the compact structures of the condensates of the N-RNA-nsp12 complex of SARS-CoV-2. The discovery of the LLPS-mediated interplay between N protein and nsp12 and the corresponding modulating compounds illuminates a feasible way to improve the accessibility of antiviral drugs (e.g., remdesivir) to their targets (e.g., nsp12/RdRp), and thus may provide useful hints for further development of effective therapeutic strategies against SARS-CoV-2.

---

To date, tens of millions of people have been infected with severe acute respiratory syndrome coronavirus 2 (SARS-CoV-2), causing the outbreak of the respiratory disease named the coronavirus disease 2019 (COVID-19). As a newly emerged member of the coronavirus family, SARS-CoV-2 is an enveloped positive-strand RNA virus, which has probably the largest genome (approximately 30 kb) among all RNA viruses. The nucleocapsid (N) protein of SARS-CoV-2 is mainly responsible for recognizing and wrapping viral RNA into helically symmetric structures, and thus insulates the viral genome from external environment^1^. It was also reported that N protein can boost the efficiency of transcription and replication of viral RNA, implying its vital and multifunctional roles in the life cycle of coronavirus^2,3^. Considering the abundant expression and high immunogenicity of N protein during viral infection, accumulating studies have been performed to investigate the potential of N protein to serve as an antiviral drug target^4-7^.

In recent years, liquid-liquid phase separation (LLPS) has been shown to offer a highly efficient mechanism to spatially and temporally modulate the cellular processes, such as signaling transduction^8,9^, transcriptional control^10-12^ and chromosome remodeling^13,14^. Currently, more and more researches have shown that LLPS also participates in multiple aspects of viral infection, including viral replication, transcriptional regulation, and the formation of stress granules to modulate host immune response^15-22^. In general, viral transcription and replication tend to take place in a specific intracellular compartment called viral factory or viral inclusion^22-24^, and particularly the replication and transcription complexes (RTCs) in coronaviruses^25-27^. Recent studies on negative-strand RNA viruses such as rabies virus, vesicular stomatitis virus (VSV) and measles virus (MeV), have indicated that LLPS is a common mechanism for N protein and phosphoprotein (P protein) to form viral inclusion-like structures^17-19^. Despite these known evidences of LLPS in negative-strand RNA viruses, it is unclear whether the N protein of SARS-CoV-2 or other coronaviruses undergoes LLPS and how the N protein driven phase separation participates in viral multiplication.

In this study, we determined the unique characteristics of the phase separation driven by the N protein of SARS-CoV-2 (termed SARS-CoV-2-N) both *in vitro* and *in vivo*. The influence of individual protein domains, mutations and nucleic acids on the phase condensation of SARS-CoV-2-N was also depicted. Importantly, the interplay between the N protein-viral RNA complex of SARS-CoV-2 (termed SARS-CoV-2-N-RNA) and nsp12, the key component of the coronavirus RTCs, was demonstrated to be mediated by LLPS. Furthermore, we found that two small molecules targeting the SARS-CoV-2-N are able to intervene the phase separation properties of the N protein-viral RNA-nsp12 (termed SARS-CoV-2-N-RNA-nsp12) complex, which may improve the accessibility of other antiviral drugs (e.g., remdesivir) to their viral targets (e.g., nsp12/RdRp). Our findings thus open a new opportunity for developing efficacious therapeutics against SARS-CoV-2.

## Results

### SARS-CoV-2-N undergoes phase separation *in vitro* and *in vivo*

The nucleocapsid (N) protein of SARS-CoV-2 (termed SARS-CoV-2-N), in common with that of other coronaviruses, contains an N-terminal RNA-binding domain (NTD) and a dimerization domain at its C terminus (CTD) (Fig. 1a and see Extended Data Fig. 1). The remaining domains of SARS-CoV-2-N are highly disordered, and contain three intrinsically disordered regions (IDRs), with one also displaying prion-like activity, according to the predictions using IUPred2^28,29^ and PLAAC^30^ programs (Fig. 1a). As accumulating evidence has shown that the proteins with intrinsically disordered and prion-like properties are generally prone to undergo liquid-liquid phase separation (LLPS)^31-34^, it is natural to ask whether SARS-CoV-2-N also displays the LLPS characteristics. To answer this question, we first expressed and purified the recombinant SARS-CoV-2-N protein with an mEGFP-tag (a monomeric variant of EGFP, A206K) or a His-tag using a prokaryotic expression system (see Extended Data Fig. 2a, b). Confocal fluorescence microscopy showed that SARS-CoV-2-N readily self-associated to form numerous micron-sized spherical condensates (Fig. 1b, c). Further time-lapse observations revealed that the SARS-CoV-2-N condensates fused and coalesced into larger ones upon their intersections (Fig. 1d, Supplementary Video 1), indicating the liquid-like properties of SARS-CoV-2-N condensates. We also used fluorescence recovery after photobleaching (FRAP) to further study the dynamics of internal molecules within the N protein condensates. Recovery of fluorescence within the bleached regions (Fig. 1e) showed that SARS-CoV-2-N can partially freely diffuse within the condensed phase, consistent with their liquid-like behavior. In addition, phase condensation of SARS-CoV-2-N was sensitive to increasing ionic strength (Fig. 1f): at low protein or high salt concentrations, SARS-CoV-2-N remained dissolved in solution and no droplets were observed, whereas at high protein or low salt concentrations, it condensed into droplets, suggesting that electrostatic interactions are important for its condensation. Collectively, the above results demonstrated that SARS-CoV-2-N is capable of undergoing LLPS *in vitro*.

**Fig.1.**
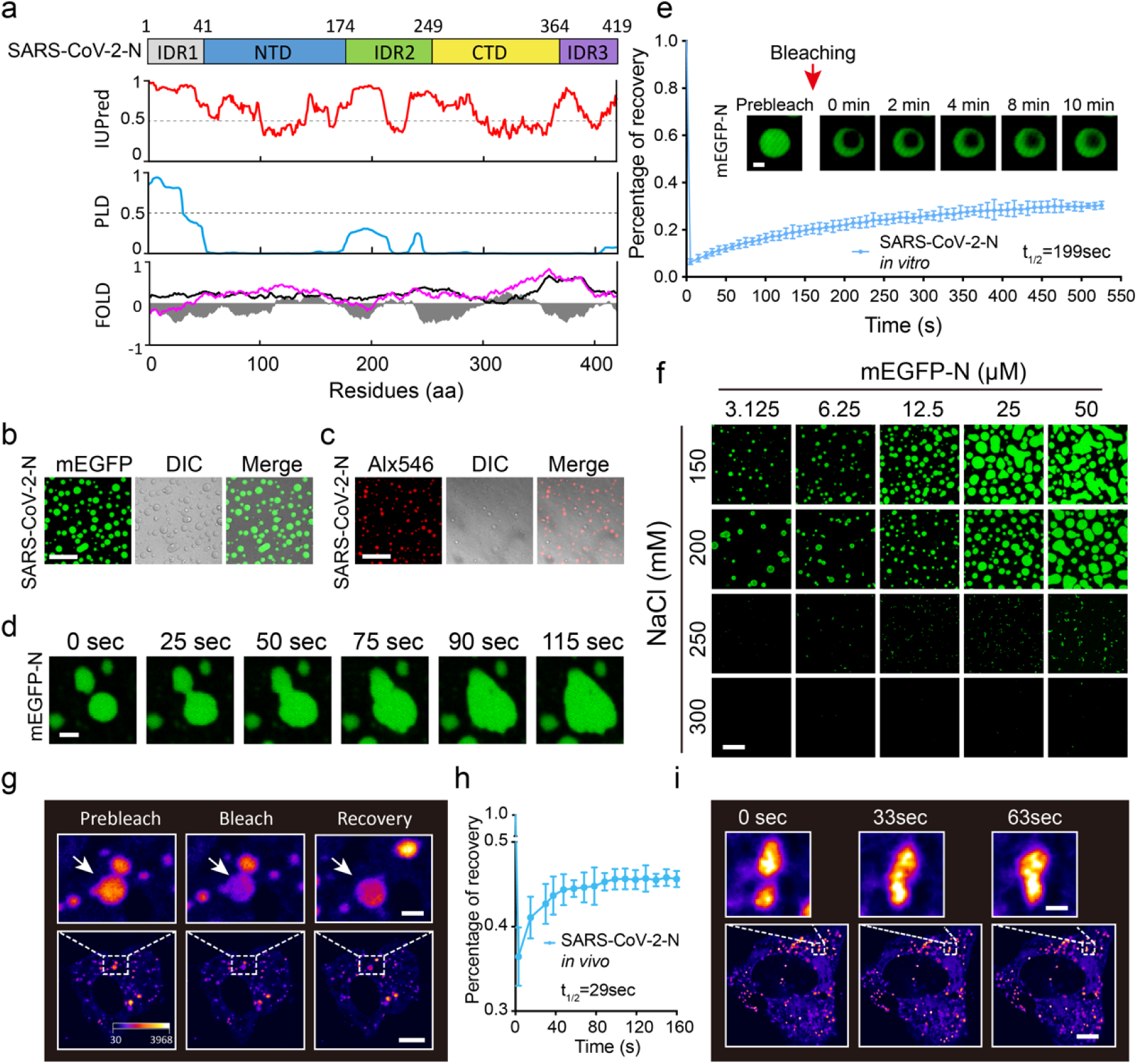
SARS-CoV-2-N undergoes phase separation *in vitro* and *in vivo*. **a**, Bioinformatic analysis of the amino acid sequence of SARS-CoV-2-N. Schematic representation of the domain structure is shown on the top. IDR, intrinsically disorder region; NTD, N-terminal domain; CTD, C-terminal domain; IUPred, prediction of intrinsic disorder; PLD, prediction of prion-like region (PLAAC); FOLD, intrinsic disorder prediction by PLAAC (purple) and prion aggregation prediction by PAPA (black). Fold index is shown in gray. **b**, *In vitro* phase separation assays of 25 μM mEGFP tagged SARS-CoV-2-N protein (mEGFP-N). Scale bar, 20 μm. **c**, *In vitro* phase separation assays of 25 μM full-length SARS-CoV-2-N protein labeled with Alx546. Scale bar, 20 μm. **d**, Fusion of mEGFP-N (50 μM) condensates. Data are representative of three independent experiments. Scale bar, 2.5 μm **e**, *In vitro* FRAP analysis of the condensates (n=3) of mEGFP-N (3 µM). Top, representative snapshots of condensates before and after bleaching. Bottom, average fluorescence recovery traces of mEGFP-N condensates. Data are representative of three independent experiments and presented as mean ± SD. Scale bar, 1 μm. **f**, Fluorescence microscopy observations of mEGFP-N condensates *in vitro* depending on different sodium chloride concentrations and protein concentrations. Scale bar, 20 µm. **g**, *In vivo* FRAP analysis of the puncta of the expressed mCherry-N (SARS-CoV-2-N tagged with mCherry) in Vero-E6 cells. Insets, representative snapshots of puncta before and after bleaching. The bleached punctum is marked with an arrow. Scale bars, 10 µm (bottom), 2 µm (insets). **h**, The average fluorescence recovery traces of the puncta (n=3) of the expressed mCherry-N protein in Vero E6 cells presented in **g**. Data are representative of three independent experiments and presented as mean ± SD. **i**, Fluorescence time-lapse microscopy of Vero E6 cells expressing mCherry-N. In **g, i**, the “Red Fire” lookup table in the NIS-Elements Viewer was used to highlight the intensity difference. Two mCherry-N puncta are zoomed-in. Scale bars, 5 μm (bottom), 1 μm (insets).

Next, we examined whether SARS-CoV-2-N can also undergo LLPS *in vivo*, by ectopically expressing an mCherry tagged version of SARS-CoV-2-N in cells. We observed that SARS-CoV-2-N formed numerous puncta-like structures upon expression in Vero E6 cells (Fig. 1g). We further evaluated the dynamicity of SARS-CoV-2-N within these puncta using FRAP, and the spatiotemporal analysis of bleaching events showed that SARS-CoV-2-N redistributed rapidly from the unbleached area to the bleached one (Fig. 1g, h). In addition, time-lapse observations revealed that the SARS-CoV-2-N puncta fused into larger condensates right after their interactions (Fig. 1i, Supplementary Video 2). All together, these results showed that SARS-CoV-2-N is also able to undergo LLPS *in vivo*.

### Viral RNA modulates phase condensation of SARS-CoV-2-N

It was previously reported that the N protein of coronavirus prefers to bind to the intergenic regions and exhibits high binding affinity with the UCUAA pentanucleotide repeats^5,35,36^. Given this fact, it would be interesting to know whether viral RNA can affect the phase separation behavior of SARS-CoV-2-N. To investigate this problem, we synthesized a 30-nt single-strand RNA of SARS-CoV-2 (genomic region 28,248-28,277), which contained a UCUAA pentanucleotide and was labeled with Cy5. We found that the addition of this viral RNA to the purified SARS-CoV-2-N solution at a physiological salt concentration (150 mM NaCl) resulted in robust co-phase separation of these two components, which was more predominant than that of a single one alone (Fig. 2a). Compared to the condensates of SARS-CoV-2-N alone (Fig. 1d), the droplets of N protein-RNA (termed SARS-CoV-2-N-RNA) complex were more spherical (Fig. 2b, Supplementary Video 3). In addition, the condensates of SARS-CoV-2-N-RNA complex exhibited slightly higher molecular exchange rates than those of N protein alone (Fig. 1e, 2c). To further study the phase separation properties of this two-component (i.e., N protein and nucleic acid) system under different conditions, we examined its condensation behaviors at different constituents. As in other two-component phase separation systems^31,37^, the complex of SARS-CoV-2-N and viral RNA also exhibited an optimal molecular ratio of condensation (Fig. 2d, e). Moreover, although the synthesized viral RNA, the host-derived RNA (β-actin RNA), the dsRNA and the ssDNA (both derived from viral RNA sequence) all induced phase separation, viral RNA displayed the most prominent effect (Fig. 2f, g) in regulating the LLPS of SARS-CoV-2-N. Thus, viral RNA plays a dominant role in promoting the phase separation of SARS-CoV-2-N *in vitro*, and modulating the liquid-like properties of the resulting condensates.

**Fig.2.**
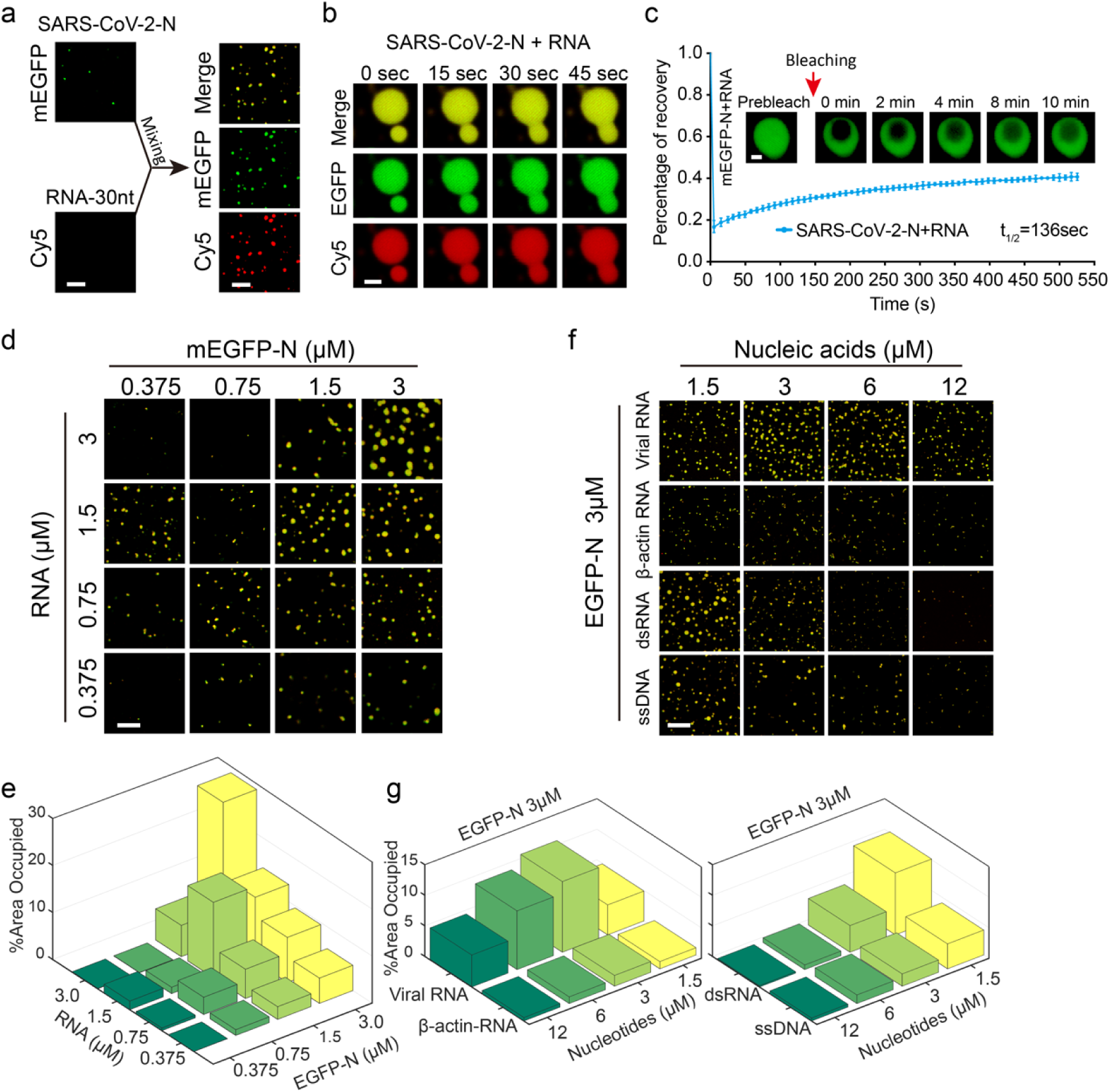
Viral RNA facilitates phase condensation of SARS-CoV-2-N. **a**, Left, *in vitro* phase separation assays of mEGFP-tagged SARS-CoV-2-N (mEGFP-N) alone (375 nM) or Cy5-labeled 30-nt viral RNA alone (1.5 μM). Right, puncta formed by mEGFP-N (375 nM) mixed with 30-nt viral RNA (375 nM) *in vitro*. Scale bar, 5 µm. **b**, Fusion upon contact of the condensates of mEGFP-N (50 μM) with 30-nt viral RNA (25 μM). Scale bar, 2.5 µm. **c**, *In vitro* FRAP analysis of the condensates (n=3) of mEGFP-N (3 µM) with 30-nt viral RNA (1.5 µM). Top, representative snapshots of condensates before and after bleaching. Bottom, average fluorescence recovery traces of mEGFP-N with viral RNA in condensates. Data are representative of three independent experiments and presented as mean ± SD. Scale bar, 1 μm. **d**, *In vitro* phase separation assays of mEGFP-N with 30-nt viral RNA at different protein/RNA concentrations. Only the merged channel is shown here. Scale bar, 10 µm. **e**, Quantitative comparison of phase condensation of mEGFP-N with 30-nt viral RNA presented in **d. f**, *In vitro* phase separation assays of mEGFP-N with nucleic acids from distinct sources and at different concentrations. Only the merged channel is shown here. Scale bar, 20 µm. **g**, Quantitative comparison of phase condensation of mEGFP-N with nucleic acids presented in **f**. In **e** and **g**, %Area Occupied = [Sum of area occupied by N protein condensates]×100/[The whole area].

### The intact structure of SARS-CoV-2-N is crucial to phase separation

To determine the contributions of individual domains of SARS-CoV-2-N to its phase separation, we first designed and expressed five truncations using a prokaryotic expression system (Fig. 3a and see Extended Data Fig. 2a) and then examined their phase separation properties at various ratios of N protein to viral RNA (Fig. 3b). For a protein/RNA ratio of 1 or 2 (i.e., 1.5 μM protein/1.5 μM RNA or 3 μM protein/1.5 μM RNA, respectively), only full-length (FL) SARS-CoV-2-N exhibited the ability to phase separate (Fig. 3b). When the protein/RNA ratio reached 4 or even 8 (i.e., 6 μM protein/1.5 μM RNA or 12 μM protein/1.5 μM RNA, respectively), although the truncations also formed phase-separated condensates, their LLPS ability was much weaker than that of the full-length SARS-CoV-2-N protein (Fig. 3b, c). All these results indicated that all domains contribute to phase separation of SARS-CoV-2-N.

**Extended Data Fig.1.**
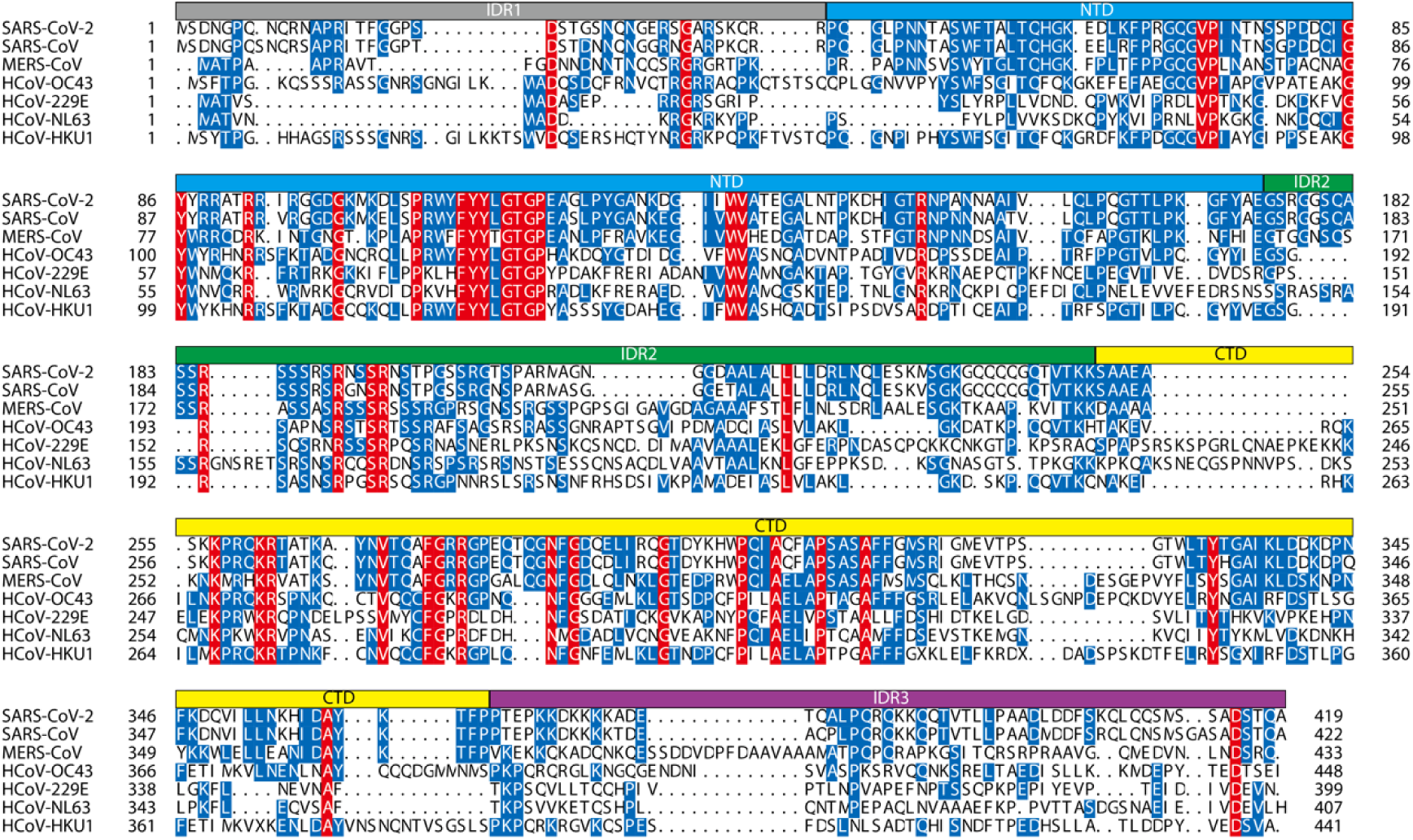
Multiple sequence alignment of the N proteins from different human coronaviruses. The nucleocapsid (N) protein sequences of the currently known coronaviruses that can infect human, including SARS-CoV-2 (GenBank: QHD43423.2), SARS-CoV (GenBank: AYV99827.1), MERS-CoV (GenBank: AVV62544.1), HCoV-OC43 (GenBank: AAR01019.1), HCoV-229E (GenBank: APD51511.1), HCoV-NL63 (NCBI Reference Sequence: YP_009328939.1) and HCoV-HKU1 (GenBank: ARU07581.1), were aligned using MUSCLE^38^. Domain architectures are depicted above the sequence alignment. The conserved residues are shaded in red, while those with the percentage of conservation larger than or equal to 50% are shaded in blue.

**Fig.3.**
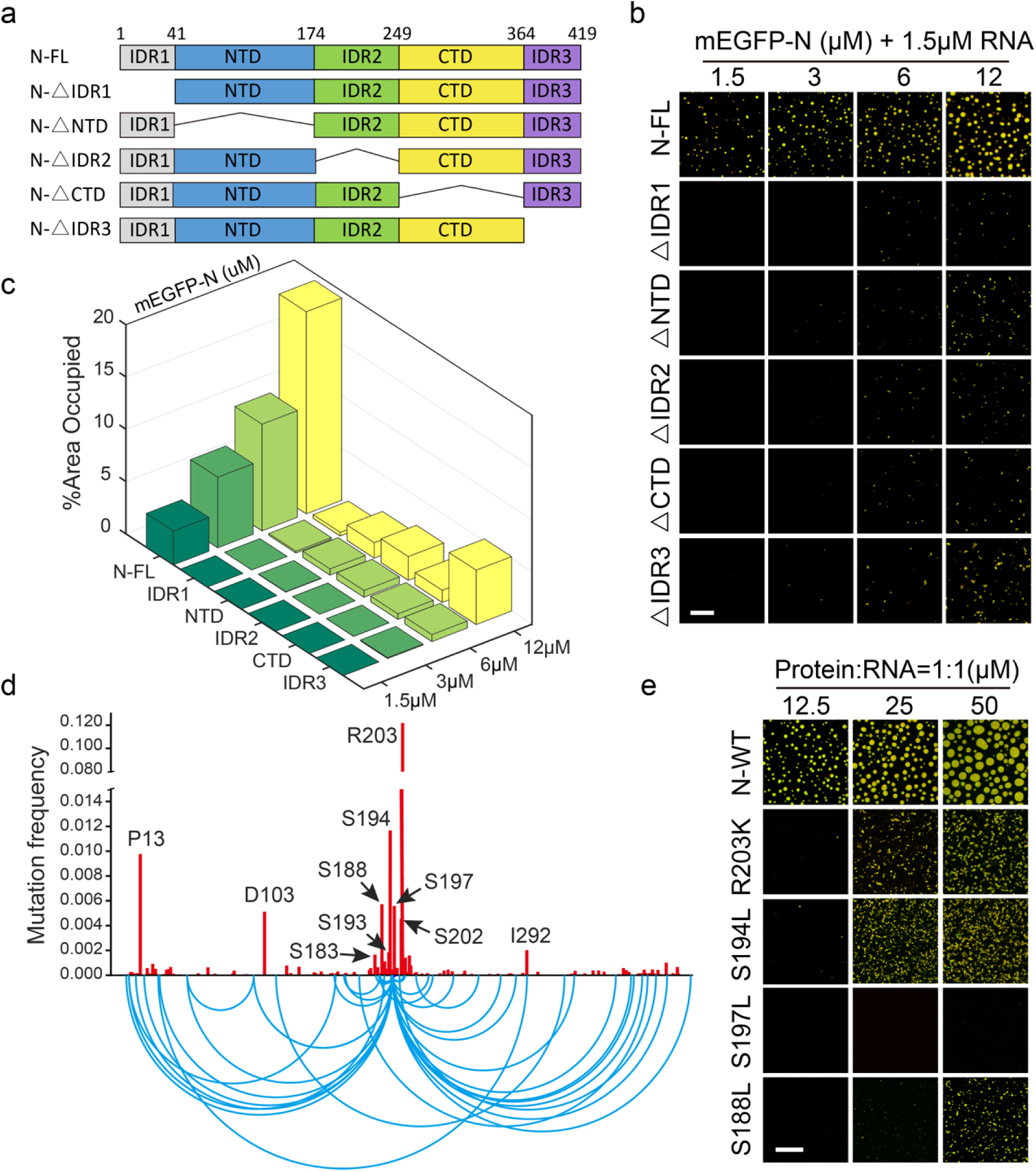
Excision of any domain and spontaneous missense mutations both compromise the LLPS of SARS-CoV-2-N-RNA complex. **a**, Diagram of the structural domains of SARS-CoV-2-N. Truncated proteins for functional analyses of different domains are shown underneath. **b**, *In vitro* phase separation assays of full-length (FL) mEGFP-N and truncations with 1.5 μM viral RNA at different concentrations of SARS-CoV-2-N (The numbers under the line represent the concentrations of the proteins). Scale bar, 20 μm. **c**, Quantitative comparison of the phase condensation of full-length and truncated mEGFP-N proteins presented in **b**. %Area Occupied = [Sum of area occupied by N protein condensates]×100/[The whole area]. **d**, Frequencies of spontaneous missense mutations in the N protein in 61,003 SARS-CoV-2 genome sequences from the China National Center for Bioinformation. Residue positions of the top 10 most frequent missense mutations are labeled. Bottom, arc diagram of double missense mutations. Only those double missense mutations with frequencies more than 0.0001 are shown. **e**, *In vitro* phase separation assays of the Alx546-labeled wild-type (WT) N protein and four mutants with mutations on the serine-arginine (SR) rich region with viral RNA of different concentrations. The ratio of N protein to viral RNA was 1:1. Scale bar, 20 μm.

**Extended Data Fig.2.**
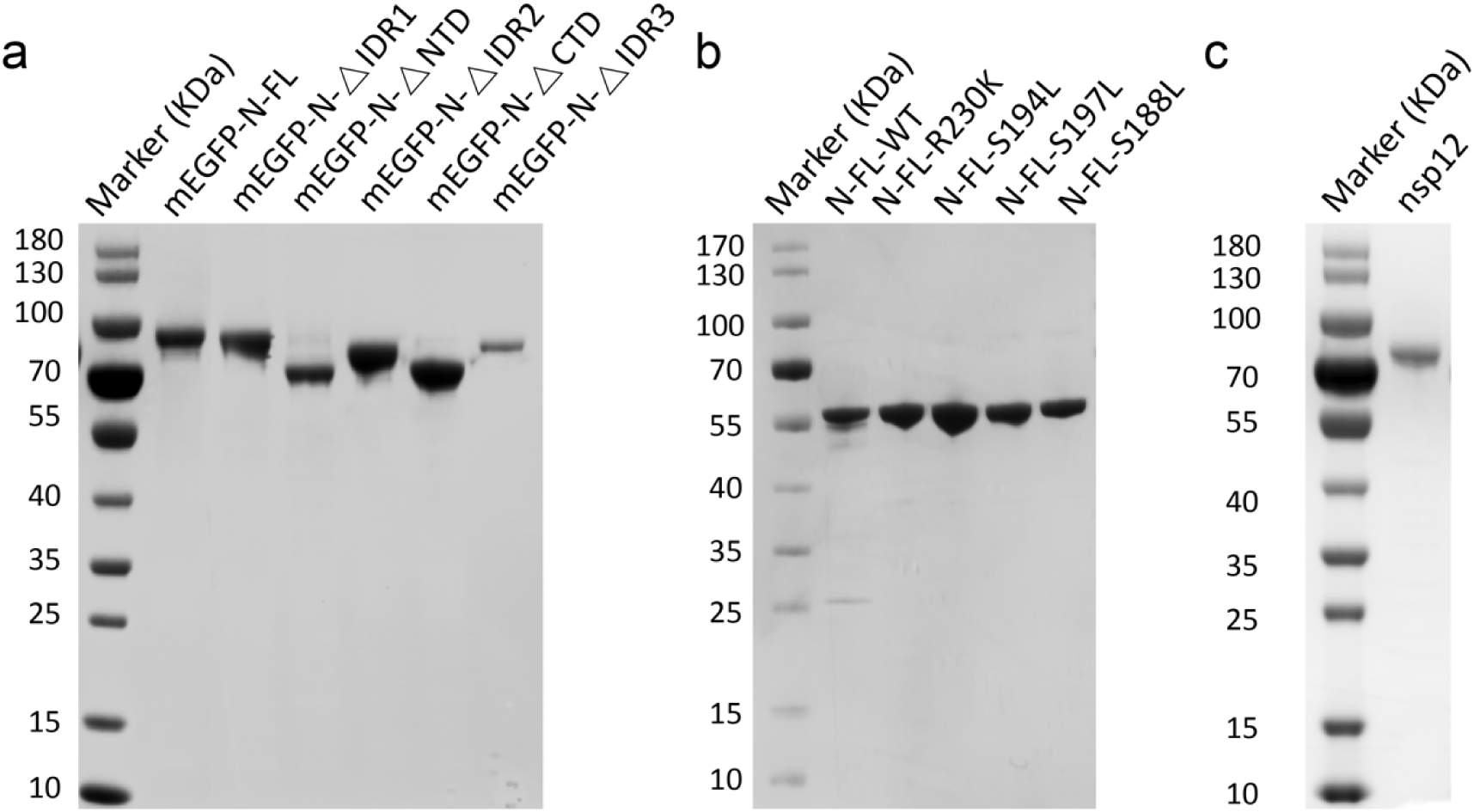
SDS-PAGE of the purified recombinant proteins of SARS-CoV-2-N used in *in vitro* assays. **a**, The mEGFP-tagged full-length (FL) and truncated proteins of SARS-CoV-2-N. **b**, The wild-type and mutant proteins of SARS-CoV-2-N with His-tagged at the N terminus. **c**. The nsp12 protein of SARS-CoV-2. The gel was stained with Coomassie Brilliant Blue.

### Spontaneous mutations impair phase separation of SARS-CoV-2-N

Since the first genome was sequenced in January 2020, 61,003 genome sequences of SARS-CoV-2 have been deposited into the China National Center for Bioinformation, 2019 Novel Coronavirus Resource^39^ (https://bigd.big.ac.cn/ncov?lang=en, July 6th, 2020). Based on these available viral genomic data, although a large number of mutations have been discovered^40,41^, their influence on virulence and pathogenicity of SARS-CoV-2 still remains largely unknown. To study the effects of mutations on the LLPS of SARS-CoV-2-N, we first collected all the missense mutations within the N protein region (between genome positions 28,274 and 29,530) (Fig. 3d). Notably, the serine-arginine (SR) rich region in the IDR2 domain of SARS-CoV-2-N was the hot spot, and harbored 7 of the top 10 most frequent mutations. The SR residues are generally considered potential phosphorylation sites, and the alternation of their phosphorylation states can impact both RNA binding and intracellular transportation of the N protein of coronavirus^42-45^. Among all the known missense mutations in SARS-CoV-2-N, R203K was of the highest frequency and always associated with other mutations (Fig. 3d), indicating that probably it was one of the earliest mutations. Next, we expressed and purified a number of mutated SARS-CoV-2-N protein *in vitro*, including R203K, S194L, S197L, and S188L mutants (4 among the top 5 mutations) (see Extended Data Fig. 2b), and then examined their phase separation capacity. Interestingly, these mutants displayed markedly weaker phase separation than the wild-type N protein (Fig. 3e). Thus, it is conceivable that the SR rich region acts as a regulatory hub to modulate the biological functions of SARS-CoV-2-N through turning its phase separation properties.

### CVL218 and PJ34 interact with SARS-CoV-2-N and affect the internal dynamics of its condensates

Considering the phase separation properties of SARS-CoV-2-N *in vitro* and *in vivo*, we want to know whether there exist small molecules or drugs that can intervene the viral life cycle through changing the condensation of this protein. According to our previous study, two poly ADP-ribose polymerase (PARP) inhibitors, CVL218 and PJ34, exhibit binding potential to SARS-CoV-2-N, as discovered by a machine learning based drug repositioning strategy^6^. Our surface plasmon resonance (SPR) assays confirmed that these two small molecules both interact with SARS-CoV-2-N, with CVL218 showing a higher binding affinity (*K*_D_=4.7 μM, Fig.4a) than PJ34 (*K*_D_=696 μM, Fig. 4b). Since the N-terminal domain (NTD) and C-terminal domain (CTD) of SARS-CoV-2-N are highly conserved and play important roles in RNA binding and self-dimerization^46^, we next investigated whether they are responsible for the interactions with these two small molecules. Unexpectedly, neither NTD nor CTD alone bound to CVL218 or PJ34 according to our *in vitro* SPR results (see Extended Data Fig. 3), implying that the entire structure and conformation of SARS-CoV-2-N, including NTD, CTD and IDRs, are essential for the interactions with CVL218 or PJ34.

**Fig.4.**
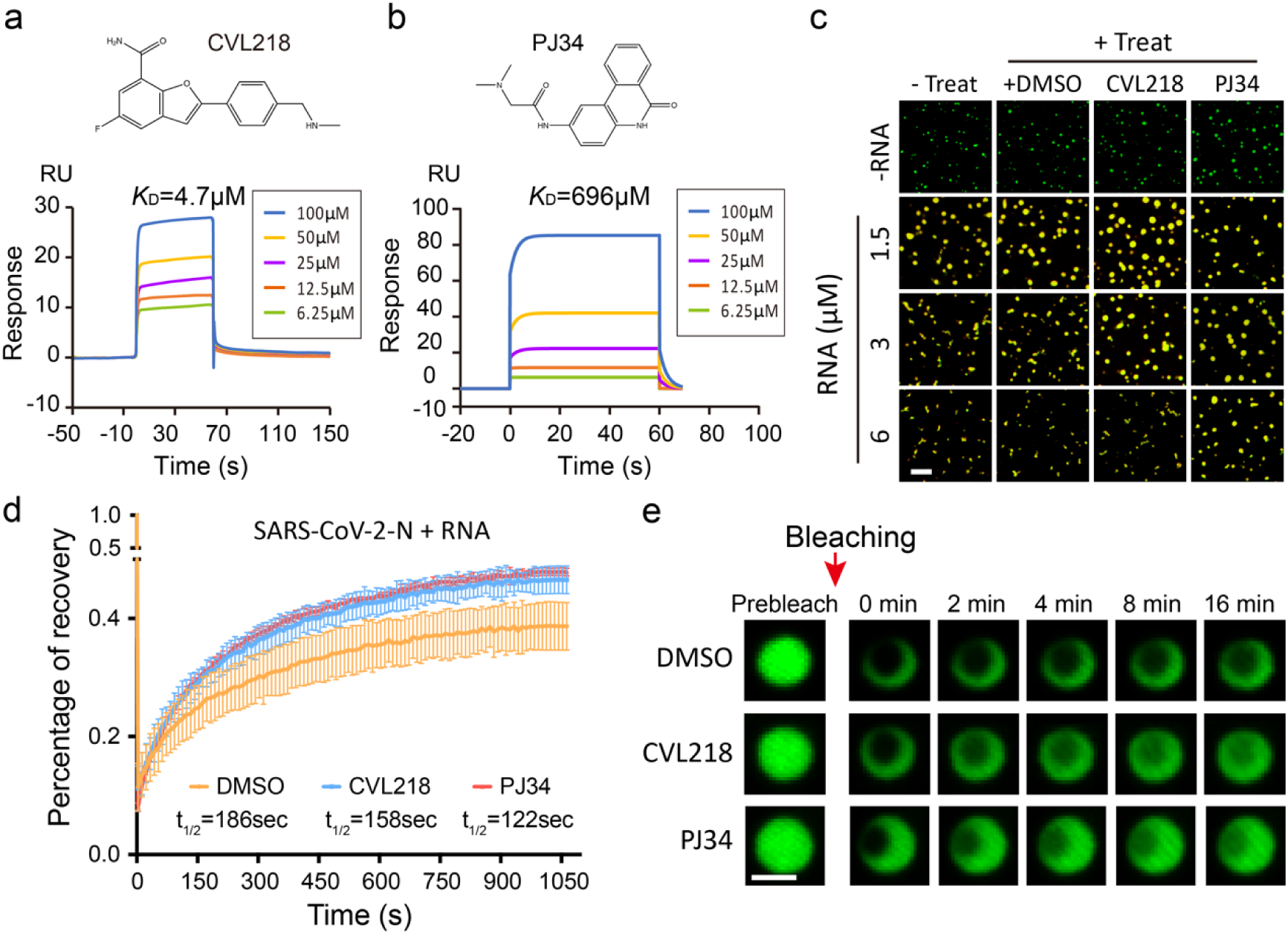
CVL218 and PJ34 bind to SARS-CoV-2-N and increase the internal dynamics of its condensates. **a, b**, Surface plasmon resonance (SPR) assays of CVL218 (**a**) and PJ34 (**b**) to the immobilized full-length SARS-CoV-2-N. Top, the chemical structures of CVL218 and PJ34, respectively. Bottom, SPR binding curves of SARS-CoV-2-N to CVL218 and PJ34, respectively. **c**, *In vitro* phase separation assays of 3 μM mEGFP-N (SARS-CoV-2-N tagged with mEGFP) with viral RNA of different concentrations and in the presence of 20 μM DMSO, CVL218 and PJ34, respectively. Only the merged channel is shown here. Scale bar, 10 μm. **d**, *In vitro* FRAP analysis of droplets (n=3) formed by mEGFP-N protein with viral RNA (mEGFP-N, 3 μM; RNA, 3 μM) under the treatment of 20 μM DMSO, CVL218 and PJ34, respectively. Data are representative of three independent experiments and presented as mean ± SD. **e**, Representative snapshots of condensates before and after bleaching presented in **d**. Scale bar, 2 μm.

Next, we examined whether CVL218 and PJ34 can influence the phase separation behavior of SARS-CoV-2-N with viral RNA. We observed that the addition of CVL218 or PJ34 had little effect on the number or morphology of the puncta of SARS-CoV-2-N-RNA complex, comparing to DMSO treatment (Fig. 4c). Nevertheless, the fluorescence intensity of the condensates with CVL218 or PJ34 treatment recovered faster than that with DMSO treatment after photobleaching (Fig. 4d, e), demonstrating that CVL218 and PJ34 can both enhance the internal mobility of the condensates of SARS-CoV-2-N-RNA complex.

**Extended Data Fig.3.**
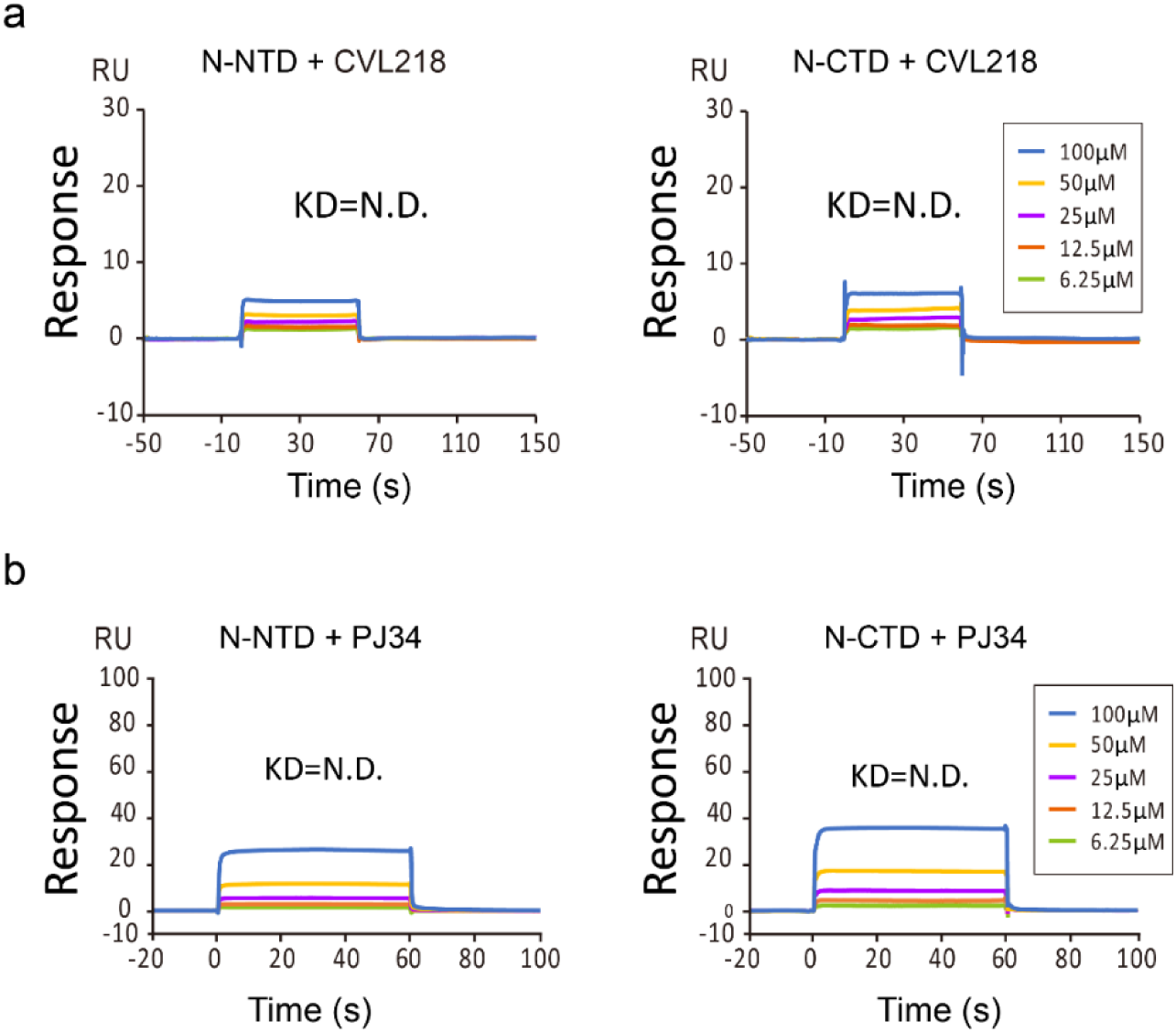
CLV218 and PJ34 do not directly bind to the NTD and CTD of SARS-CoV-2-N. **a, b**, Surface plasmon resonance (SPR) assays of CVL218 (**a**) and PJ34 (**b**) to the immobilized N-terminal domain (NTD) and C-terminal domain (CTD) of SARS-CoV-2-N.

### Nsp12 can be recruited into the SARS-CoV-2-N-RNA condensates

Nsp12, a core component of RNA dependent RNA polymerase (RdRp) complex in SARS-CoV-2, forms the coronavirus transcription and replication machinery with nsp7 and nsp8 together (Fig. 5a). It was previously reported that N protein can cooperate with RdRp to facilitate viral infection^25^. However, the underlying mechanisms still remain largely obscure. To further explore whether the interplay between N protein and RdRp of SARS-CoV-2 is mediated by phase separation, we also purified the nsp12 protein (see Extended Data Fig. 2c) and examined its potential involvement in LLPS. We observed that nsp12 alone cannot undergo phase separation under physiological salt condition (Fig. 5b), whereas it readily converted to amorphous condensates when mixed with viral RNA (see Extended Data Fig. 4). In addition, FRAP assays indicated that the fluorescence intensity of nsp12-RNA condensates cannot recover after photobleaching (Supplementary Video 4), implying that little molecular exchange occurs within the resulting solid-state condensates. Nevertheless, we found that nsp12 can be readily recruited into the SARS-CoV-2-N-RNA condensates without changing their morphological shapes and arrangements (Fig. 5b). Here, the highly concentrated condensates of SARS-CoV-2-N-nsp12 complex may provide a favorable condition for fast viral replication *in vivo*.

**Extended Data Fig.4.**
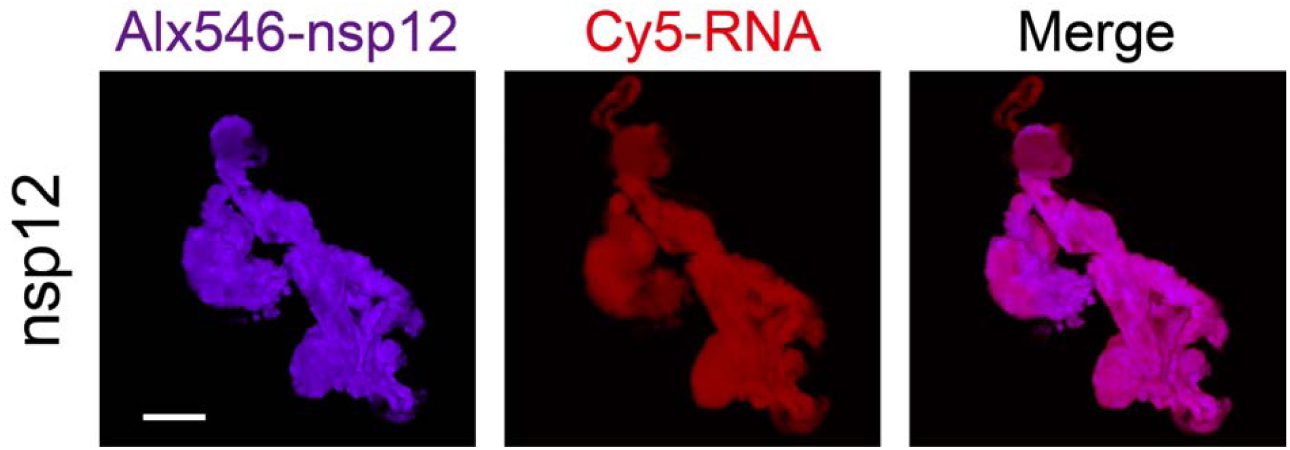
Nsp12 and viral RNA of SARS-CoV-2 form amorphous condensates *in vitro*. Fluorescence microscopy images of 3 μM Alx546-labeled nsp12 (purple) mixed with 3 μM Cy5-labeled viral RNA (red) of SARS-CoV-2. Scale bar, 5 μm.

**Fig.5.**
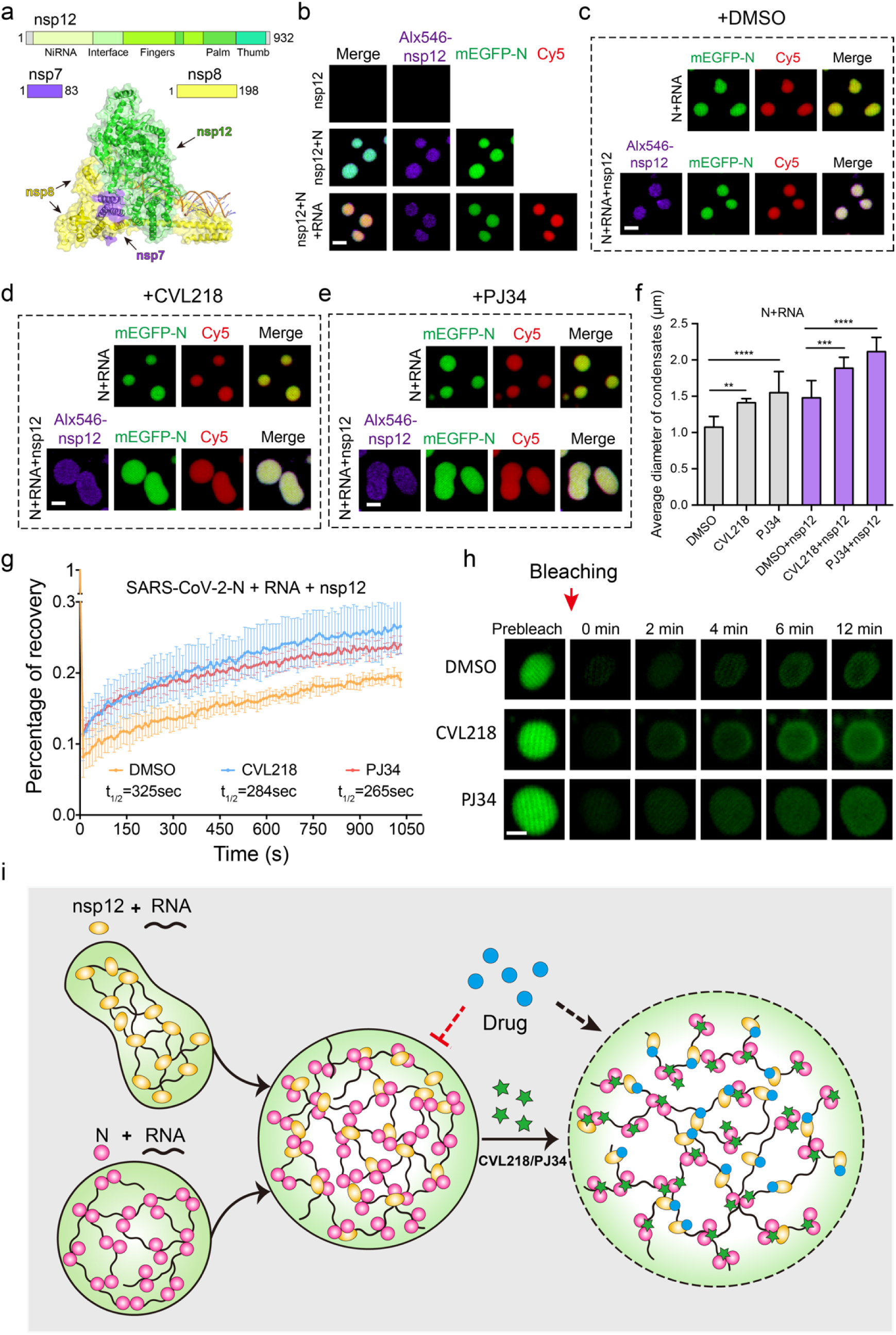
CVL218 and PJ34 influence the morphology and internal dynamics of the condensates of SARS-CoV-2-N-RNA-nsp12 complex. **a**, Schematic diagram of the structure of RNA-dependent RNA polymerase (RdRp) complex. Top, domain architectures of nsp12, nsp7 and nsp8, respectively. Bottom, a ribbon and surface view of the overall structure of RdRp complex (PDB code: 6yyt). **b**, *In vitro* phase separation assays for 3 μM Alex546 labeled nsp12 (purple) alone, with 3 μM mEGFP-N (SARS-CoV-2-N tagged with mEGFP, green) and with the complex of 3 μM mEGFP-N and 3 μM Cy5-labeled viral RNA (red), respectively. The molar ratio between mEGFP-N and RNA is 1:1. Scale bar, 2 μm. **c-e**, *In vitro* phase separation assays of 3 μM mEGFP-N protein with 3 μM viral RNA in the absence and presence of 3 μM nsp12 under the treatment of 20 μM DMSO (**c**), CVL218 (**d**) and PJ34 (**e**), respectively. Scale bar, 2 μm. **f**, Quantification of the effect of CVL218 or PJ34 treatment on the average diameters of the condensates presented in **c-e**. The diameters were measured from the fluorescence microscopy images and shown as mean±SD over three independent experiments. P values were determined by one-way analysis of variance (ANOVA) with Tukey’s multiple comparison test, **: P < 0.01, ***: P < 0.001, ****: P < 0.0001. **g**, *In vitro* FRAP analysis of the condensates (n=3) of SARS-CoV-2-N-RNA-nsp12 complex (mEGFP-N, 3 μM; RNA, 3 μM; nsp12, 3 μM) under the treatment of 20 μM DMSO, CVL218 and PJ34, respectively. Data are representative of three independent experiments and presented as mean ± SD. **h**, Representative snapshots of the condensates before and after bleaching presented in **g**. Scale bar, 2 μm. **i**, A model mechanism of the inhibition of viral replication and transcription of SARS-CoV-2 by small molecules in a phase separation dependent manner. Nsp12 alone cannot undergo phase separation *in vitro*, but it can be recruited into the droplets of SARS-CoV-2-N-RNA complex, despite the fact that nsp12 and viral RNA can form solid-state condensates (see Extended Data Fig. 5). Comparing to those with DMSO treatment, the diameters and mobility of SARS-CoV-2-N-RNA-nsp12 droplets obviously increased after the treatment of CVL218/PJ34, which can attenuate the local density of the condensates and thus promote the entrance of other antiviral drugs (e.g., remdesivir) into their targets (e.g., nsp12/RdRp).

### CVL218 and PJ34 affect the morphology and condensation properties of SARS-CoV-2-N-RNA-nsp12 complex

Next, we sought to examined whether CVL218 or PJ34 can affect the phase condensation properties of SARS-CoV-2-N-RNA-nsp12 complex. Interestingly, the sizes of SARS-CoV-2-N-RNA-nsp12 condensates in the CVL218 or PJ34 treated group were much larger than those of the corresponding group without nsp12 (Fig. 5d-e and see Extended Data Fig. 5). More importantly, no matter with or without nsp12, the sizes of SARS-CoV-2-N-RNA condensates under CVL218 or PJ34 treatment were significantly larger than those of the DMSO treated group (Fig. 5c-f). Moreover, FRAP assays indicated that the fluorescence recovery rates of SARS-CoV-2-N-RNA-nsp12 condensates were faster in the CVL218 or PJ34 treated group than those of the control treatment (Fig. 5g, h). Thus, our results suggested that CVL218 and PJ34 can induce a more dynamic condition to facilitate the intermolecular exchange of internal molecules within the droplets of SARS-CoV-2-N-RNA-nsp12 complex *in vitro*. The accelerating exchange rates of internal molecules within the droplets may loosen the solid-state condensation of the nsp12-containing RdRp complex and thus benefit the entrance of other antiviral drugs (e.g., remdesivir) into their targets (e.g., nsp12/RdRp).

**Extended Data Fig.5.**
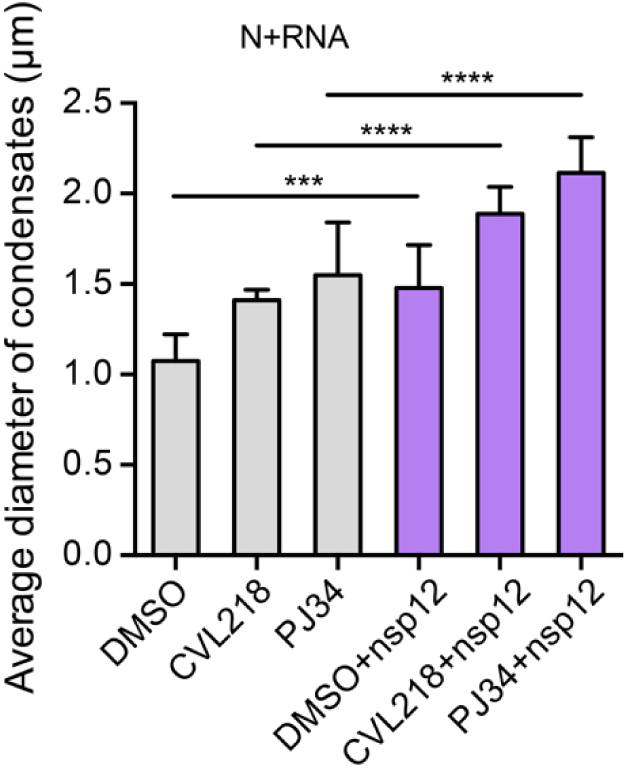
The recruitment of nsp12 enlarges the sizes of the SARS-CoV-2-N-RNA condensates. Supplementary quantitative comparison of average diameters of the condensates in the presence and absence of nsp12 presented in Fig.5 c, d, e. The diameters were measured from the fluorescence microscopy images and shown as mean±SD over three independent experiments. P values were determined by one-way analysis of variance (ANOVA) with Tukey’s multiple comparison test, ***: P < 0.001, ****: P < 0.0001.

**Extended Data Fig.6.**
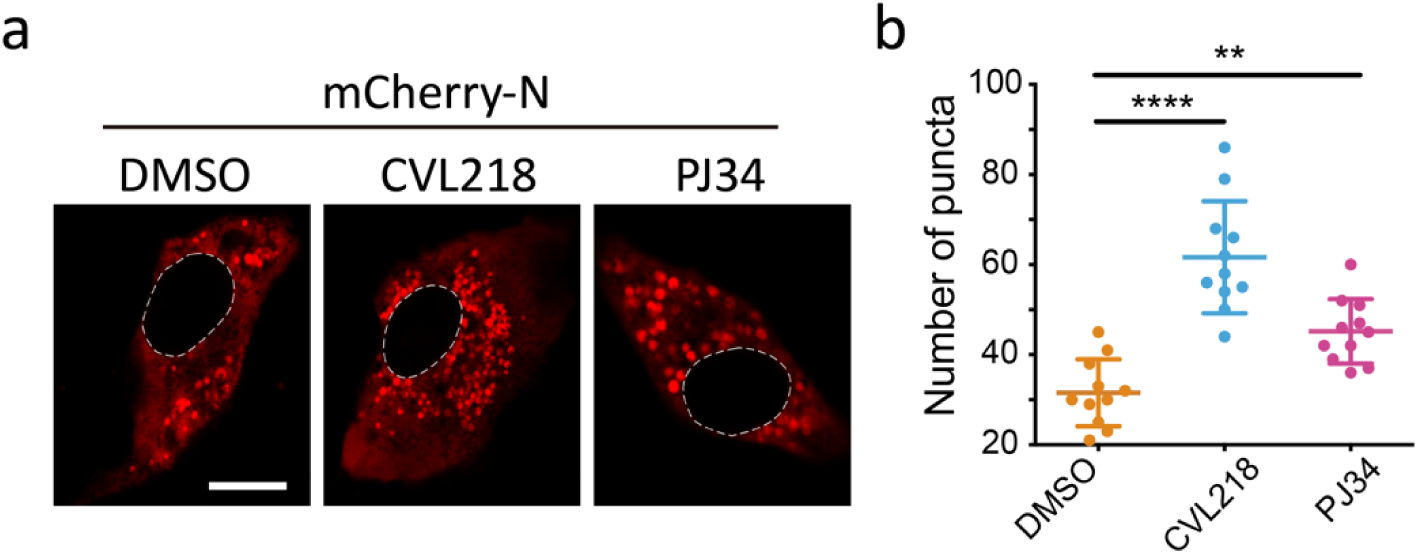
CVL218 and PJ34 treatments influence the condensation of SARS-CoV-2-N *in vivo*. **a**, Droplet formation of the expressed mCherry-N (SARS-CoV-2-N tagged with mCherry) in Vero-E6 cells for 48h. Scale bar, 10 μm. **b**, Quantitative comparison of the numbers of droplets presented in **a**. In total 11 transfected cells were considered in each group (i.e., DMSO, CVL218-treated or PJ34-treated). Data are shown as mean±SD and P values were determined by one-way analysis of variance (ANOVA) with Dunnett’s multiple comparison test, **: P < 0.01, ****: P < 0.0001.

## Discussion

Our results showed that the N protein of SARS-CoV-2 is capable of undergoing LLPS, and extra addition of viral RNA further facilitates its phase separation, which has also been recently confirmed by several independent research teams^47-49^. Moreover, with in-depth study, we discovered that nsp12/RdRp can be recruited into the condensates of SARS-CoV-2-N-RNA complex. Furthermore, in terms of the intervention of the SARS-CoV-2-N driven phase separation, we set up an experimental workflow for compound verification and proposed a conceptual framework for developing the small molecule-based therapeutics against SARS-CoV-2, based on a rationale of combination therapy to target both N protein and nsp12/RdRp.

Our work suggests that LLPS is likely to be the driving force of the connection between N protein and the replication and transcription complexes (RTCs), and thus altering the N-driven LLPS can possibly intervene viral multiplication and infection. Normally, many non-structural proteins (nsps) of coronavirus anchor in double-membrane vesicles and convoluted membranes and are packaged into RTCs in infected cells^25-27^. The dynamics of RTCs is generally relatively low, implying that they display relatively static structures and probably their communications with the surroundings rely on other components^26^. Yet, N protein is so far the only known structural protein of coronavirus that shuttles into and outside RTCs and plays a vital role in coordination with the RdRp complex^25-27^. Likewise, in the negative-strand RNA viruses, a granular structure termed inclusion body particularly stays in cytoplasm serving as a site of viral replication and transcription^15,17-19,22^. Interestingly, two structural proteins, i.e., N and P proteins, have been shown to be sufficient to spontaneously form an inclusion body-like structure mediated by LLPS in rabies virus, vesicular stomatitis virus (VSV) and measles virus (MeV)^17-19^. Here, we showed that in SARS-CoV-2, nsp12 can be recruited into the condensates of N protein, which are then turned to exhibit more liquid-like characteristics. Therefore, LLPS is probably an efficient mechanism to organize N protein and nsp12 together in SARS-CoV-2 to achieve fast viral replication. Of course, more evidence is needed to answer whether LLPS is a common mechanism for the formation of viral-specific compartments in other viruses.

Although extensive studies have been conducted on developing effective therapeutics against coronavirus through primarily targeting the spike protein and viral enzymes (e.g., nsp12/RdRp, 3C-like protease and papain-like protease), there is little success of these strategies. According to our results, we speculate that the LLPS characteristics of N protein may be one of the underlying reasons for the failure of many anti-coronavirus drugs. A related evidence comes from a recent study showing that a number of antineoplastic drugs cannot freely diffuse, but rather become partitioned in specific protein condensates *in vitro*^50^. This result supports the hypothesis that altering the biophysical properties of condensates may improve the distribution and efficacy of drugs. In our study, we observed that *in vitro* nsp12 and viral RNA form amorphous condensates, implying that their complex is relatively immobile, which is also consistent with the previous observations on the dynamics of RTCs^25-27^. The solid state of nsp12-RNA complex may exclude the nsp12 targeting drugs (e.g., remdesivir^51-53^) from the surrounding solution, which thus provides a possible explanation on why many nucleotide analog drugs as broad-spectrum viral RdRp inhibitors have poor performance in the treatment of coronavirus^54,55^.

As a proof of concept on discovering an intervention of the N protein driven LLPS, we identified two small molecules, i.e., CVL218 and PJ34, with the ability to affect the condensation properties of the SARS-CoV-2-N-nsp12 complex. These two compounds can influence the morphology of the N protein driven condensates, and also augment their sizes *in vitro* (Fig. 5f). Meanwhile, we also observed an increasing number of puncta of overexpressed mCherry-N in Vero-E6 cells, after CVL218/PJ34 treatment (see Extended Data Fig. 6a, b). In addition, these two small molecules both tune the SARS-CoV-2-N-nsp12 droplets to become more liquid-like, thus increasing the exchange with the surrounding solution and attenuating the molecular interactions within the condensates. Based on these observations, we speculate that CVL218 and PJ34 may act as bulking agents to reduce the local density of the SARS-CoV-2-N-nsp12 condensates. The increasing penetrability may possibly contribute to the access of other small-molecule drugs into the condensates. Therefore, we propose that CVL218 or PJ34 can be applied to facilitate the entrance of other antiviral drugs (e.g., remdesivir^51-53^) into RTCs by remodeling the communications between N protein and RTCs and promoting the permeability of RTCs (Fig. 5i). Our results suggest that the N protein driven LLPS is a promising target for the design of antiviral drugs, and the deep understanding into the functional roles of N protein in regulating the accessibility of RTCs will thus advance the development of anti-SARS-CoV-2 therapies.

## Methods

### Cell culture and transfection

Vero E6 cells were kindly provided by Dr. Yiyue Ge and Dr. Jingxin Li from NHC Key Laboratory of Enteric Pathogenic Microbiology, Jiangsu Provincial Center for Disease Control and Prevention. Vero E6 cells were cultured in Dulbecco’s modified Eagle’s medium (HyClone) supplemented with 10% fetal bovine serum (HyClone SH30071.03 and SH30396.03) and maintained at 37°C in a humidified incubator with 5% CO_2_. FuGENE 6 (Promega, E2691) was used for transient transfection according to the manufacturer’s instructions.

### Construction of recombinant plasmids

The recombinant plasmids of pET28a-N and pET22b-nsp12 were kindly provided by Cellregen Co., Ltd. and Prof. Zhiyong Lou^56^, respectively. The mutants of pET28a-N were constructed by seamless cloning kits (Beyotime, D7010M) and confirmed by sequencing. For the construction of mEGFP-N plasmids, the full-length gene and truncations of SARS-CoV-2-N were both cloned into a PL118 vector (an in-house modified vector based on pRSFDuet1) containing an N-terminal 6×his-mEGFP tag, respectively. The full-length gene of SARS-CoV-2-N was cloned into a pCDNA3.1 vector containing an N-terminal mCherry tag. The detailed primer sequences are listed in Supplementary Table 1.

### Protein expression and purification

The recombinant full-length mEGFP-N protein and truncations were overexpressed in *E. coli* BL21 (DE3). After overnight induction by 0.2mM isopropyl β-D-thiogalactoside (IPTG) at 16 °C in LB medium, cells were harvested and suspended in the buffer (40mM HEPES, pH 7.5, 1M NaCl, 20mM imidazole and 2mM phenylmethylsulfonyl fluoride). After cell lysis and centrifugation, the recombined proteins were purified to homogeneity over HisTrap column and eluted with a linear imidazole gradient from 20 mM to 500 mM. The proteins were further purified by size-exclusion chromatography using a Superdex 200 Increase 10/300 GL column (GE Healthcare) in elution buffer (40 mM HEPES, pH7.5, 1M NaCl, 5% glycerol, 1 mM EGTA, 1 mM MgCl_2_). The purification procedures of the recombinant wild-type pET28a-N protein and mutants were essentially the same as that of the mEGFP-N protein except for a different size-exclusion chromatography buffer (20mM Tris, pH 7.5, 300mM NaCl).

### Protein labeling

All pET28a-N proteins (WT and mutants) and nsp12 protein were labeled by incubating with a 1:1 molar ratio of Alexa Fluor™ 546 carbox (Thermo Fisher Scientific) for 1 h at room temperature with continuous stirring. Then, the free dyes were removed by centrifugation in MICROSPIN G-50 column (GE Healthcare, 27-5330-01). The labeled proteins were stored at −80°C. For *in vitro* phase separation experiments, 5% labeled protein was mixed with unlabeled before use.

### Synthesis of RNA and DNA

The 5’-Cy5-labeled 10-bp viral RNA oligo (AGCUGAUGAG) and 30-bp RNA oligos (viral RNA: GAUUUCAUCUAAACGAACAAACUAAAAUGU; human β actin RNA: UCACCAACUGGGACGACAUGGAGAAAAUCU) with were synthesized at HIPPOBIO, LLC. The double-strand RNA was annealed at 25 μM in the annealing buffer (40 mM HEPES, pH 7.4, and 150 mM NaCl) using a thermocycler, during which the oligos were heated up to 95 °C for 2 min and gradually cooled to 25°C over an hour.

### Phase separation assays

*In vitro* LLPS experiments were performed at room temperature. All samples were seeded and recorded on 384 low-binding multi-well 0.17 mm microscopy plates (In Vitro Scientific) and sealed with optically clear adhesive film. For phase separation assays with the mEGFP-N protein of SARS-CoV-2, solutions of GFP fusion proteins were diluted to the indicated final concentrations in 20 mM HEPES, pH 7.4, 500 mM NaCl, 5% glycerol, 1 mM EGTA and 1mM MgCl_2_ in a total volume of 10 µl to induce phase separation. For the N proteins without tags, the assays were performed in 20 mM Tris-HCl, pH 7.5 and 150 mM NaCl. For phase separation assays treated with small molecules, CVL218 or PJ34 (dissolved in 1% DMSO) were added to the well mixed phase separation samples prior to imaging at a final concentration of 20 μM. The group treated with 1% DMSO was used as the control.

For *in vivo* assays, Vero E6 cells were seeded into 4-well chamber 35 mm dishes with a density of 5 × 10^5^ cells/well. For cells to reach 70% confluent, 1 µg pCDNA3.1-mcherry-N plasmid was transfected, with the replacement of normal cell culture medium by that supplemented with CVL218 or PJ34 at a final concentration of 20 µM. For the control wells, cell medium containing 1% DMSO was added. Imaging was performed with a NIKON A1 microscope equipped with a 100× oil immersion objective. NIS-Elements AR Analysis was used to analyze the images.

### Fluorescence recovery after photobleaching (FRAP) measurements *in vivo* and *in vitro*

FRAP experiments were carried out with a NIKON A1 microscope equipped with a 100× oil immersion objective. Droplets were bleached with the corresponding laser pulse (3 repeats, 80% intensity, and dwell time 1 s). Recovery from photobleaching was recorded for the indicated time point.

### Mutation frequency analysis

To perform the mutation frequency analysis of SARS-CoV-2-N protein, we used 61,003 SARS-CoV-2 genome sequences downloaded from the China National Center for Bioinformation, 2019 Novel Coronavirus Resource^39^ (downloaded on July 6th, 2020). We considered all the missense mutations among the N protein region (from positions 28,274 to 29,530 in the genome).

### Surface plasmon resonance assays

Surface plasmon resonance (SPR) assays were performed on Biacore S200 with a CM5 sensor chip (GE Healthcare Life Sciences) at room temperature. The full-length N protein, NTD and CTD of SARS-CoV-2 were all diluted in 10 mM sodium acetate (pH 5.0) and immobilized on a CM5 sensor chip by amine coupling. The running buffer contained 1×PBS-P with 2% DMSO. The tested drugs (CVL218 or PJ34) in 2-fold serial dilutions were made in the running buffer. The solutions flowed through the chip surface at a flow rate of 20 μL/min at room temperature (25°C). The dissociation constants (*K*_D_) of CVL218 and PJ34 binding to full-length N protein, NTD and CTD were calculated from the association and dissociation curves of the sensorgrams using the BIA evaluation program (Biacore).

## Acknowledgements

We thank Dr. Tingting Li for helpful discussions on phase separation. We acknowledge the assistance of Protein Preparation and Identification Facility (Technology Center for Protein Science, Tsinghua University) for protein expression and purification, High Throughput Screening (HTS) Core Facility for SPR experimental guidance (Center of Pharmaceutical Technology, Tsinghua University) and SLSTU-Nikon Biological Imaging Center (Center of Pharmaceutical Technology, Tsinghua University) for imaging support. This work was supported in part by the National Natural Science Foundation of China (61872216 and 81630103 to JZ, 31900862 to DZ, 31871443 to PL), the National Key R&D Program (2019YFA0508403 to PL), the Turing AI Institute of Nanjing and the Zhongguancun Haihua Institute for Frontier Information Technology.

## Author Contributions

DZ, WX, PL and JZ conceived the research project. PL and JZ supervised the study. DZ, WX and XZ designed and performed experiments, and analyzed the data. XT, SW and XF performed protein expression and purification *in vitro*. EY, YX and PL analyzed the SARS-CoV-2 genomic data. NW and HS analyzed the structural data of SARS-CoV-2-N. XS supplied PJ34 and CVL218 and provided discussions on the SPR results. HL participated in the project discussion and provided suggestions. DZ, WX, XZ, PL and JZ wrote the manuscript with support from all authors.

