## Supplementary figures and images for "Understanding the phase separation characteristics of nucleocapsid protein provides a new therapeutic opportunity against SARS-CoV-2"

### Extended Data Fig1.png

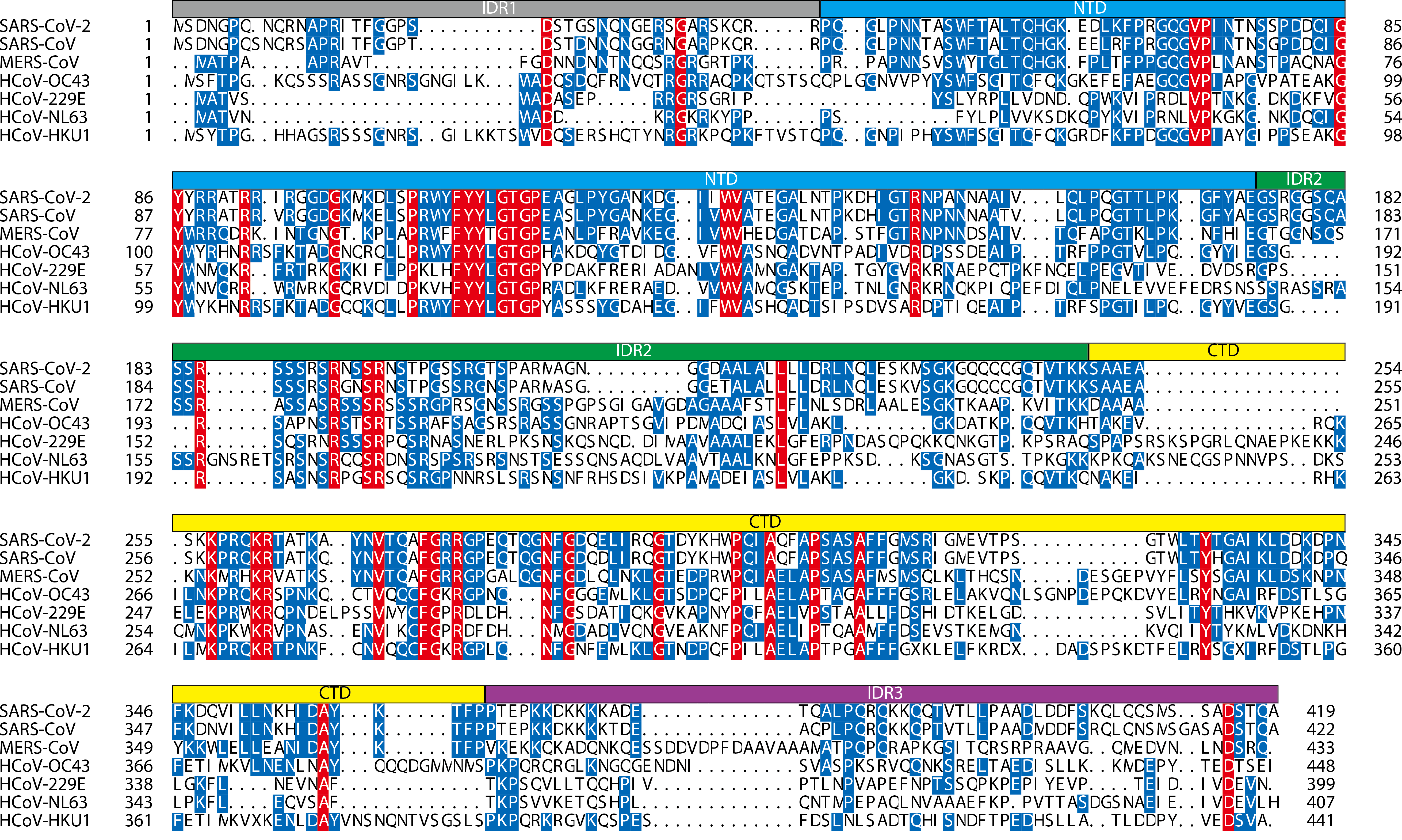

### Extended Data Fig2.png

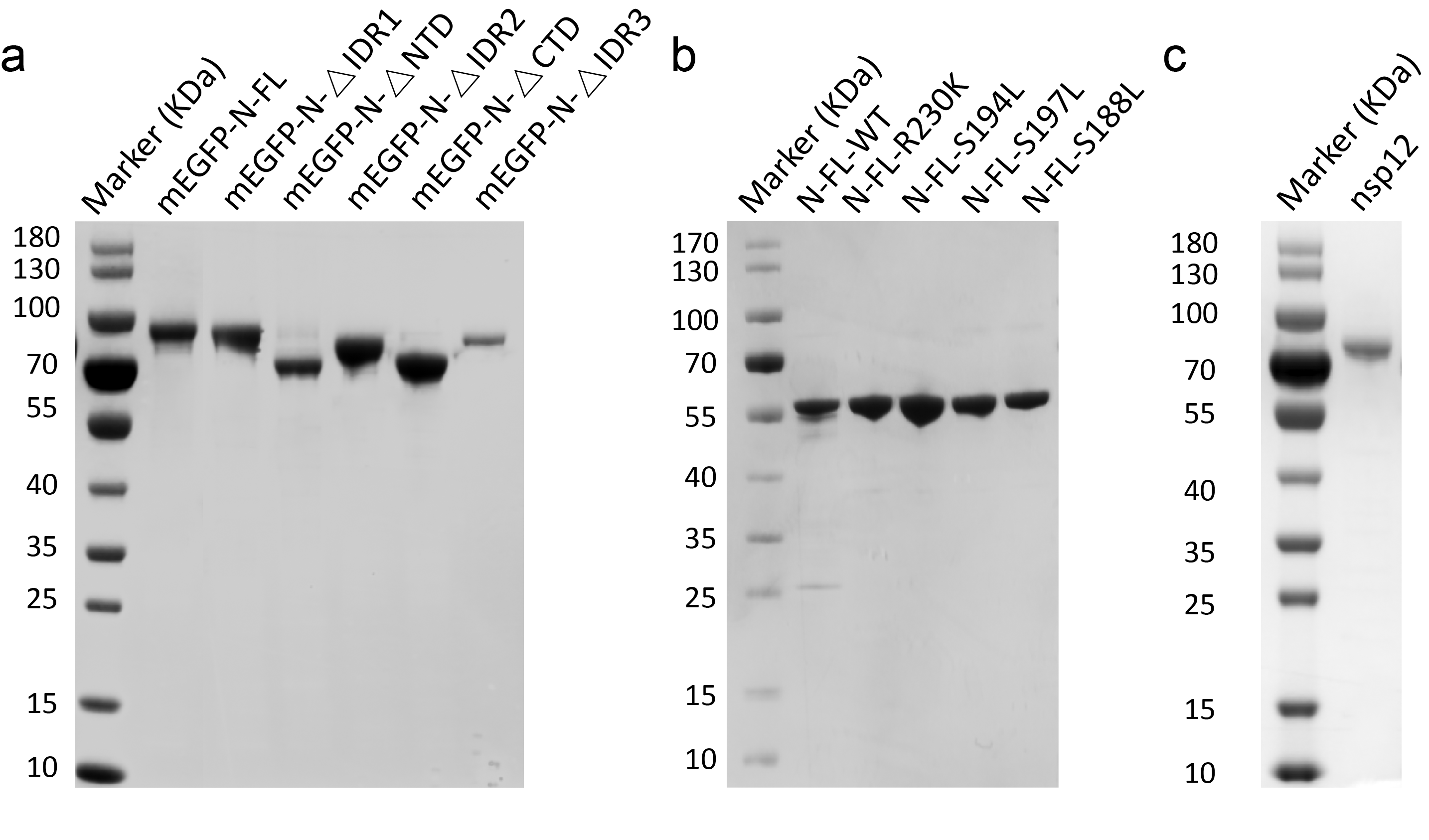

### Extended Data Fig3.png

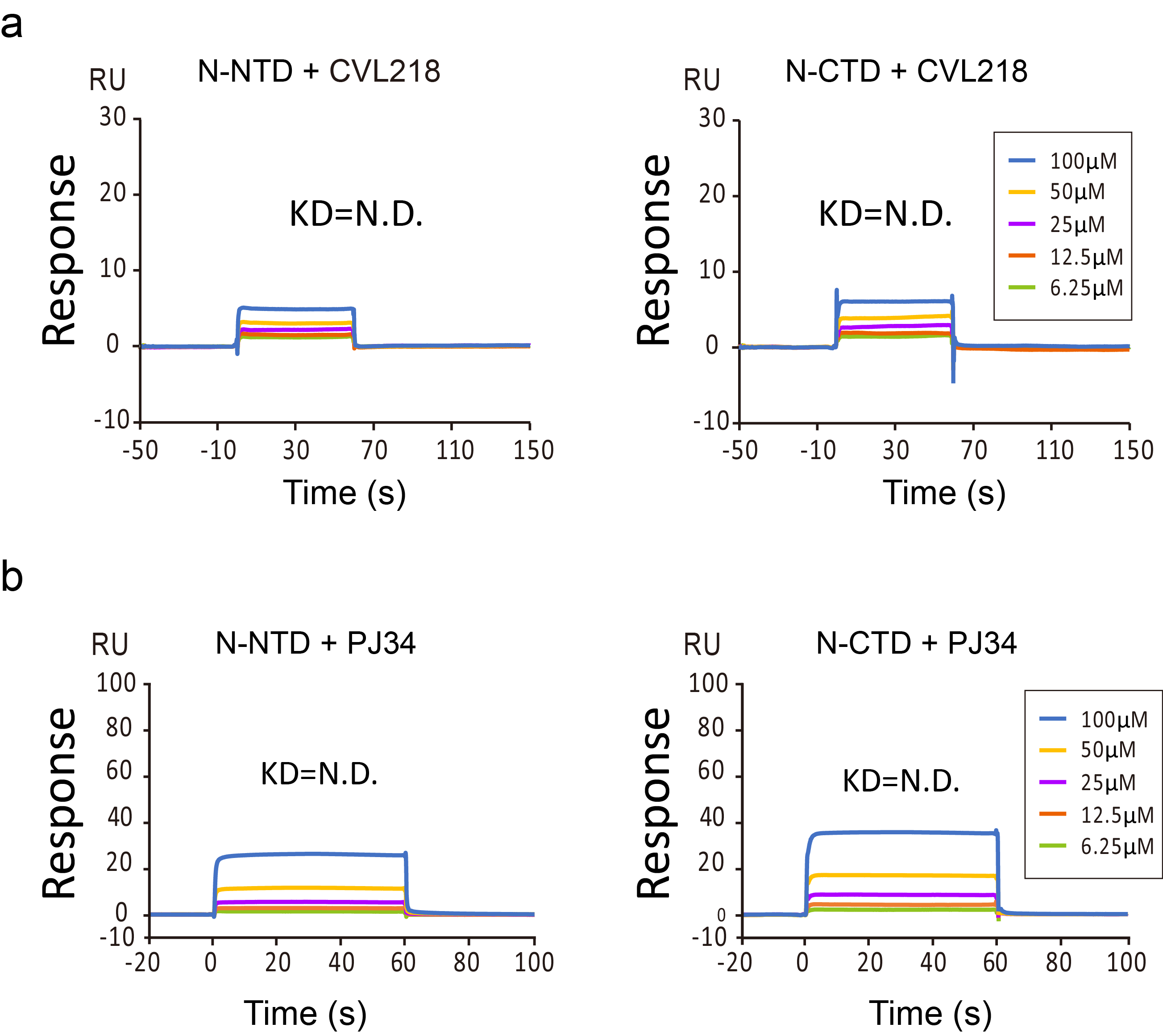

### Extended Data Fig4.png

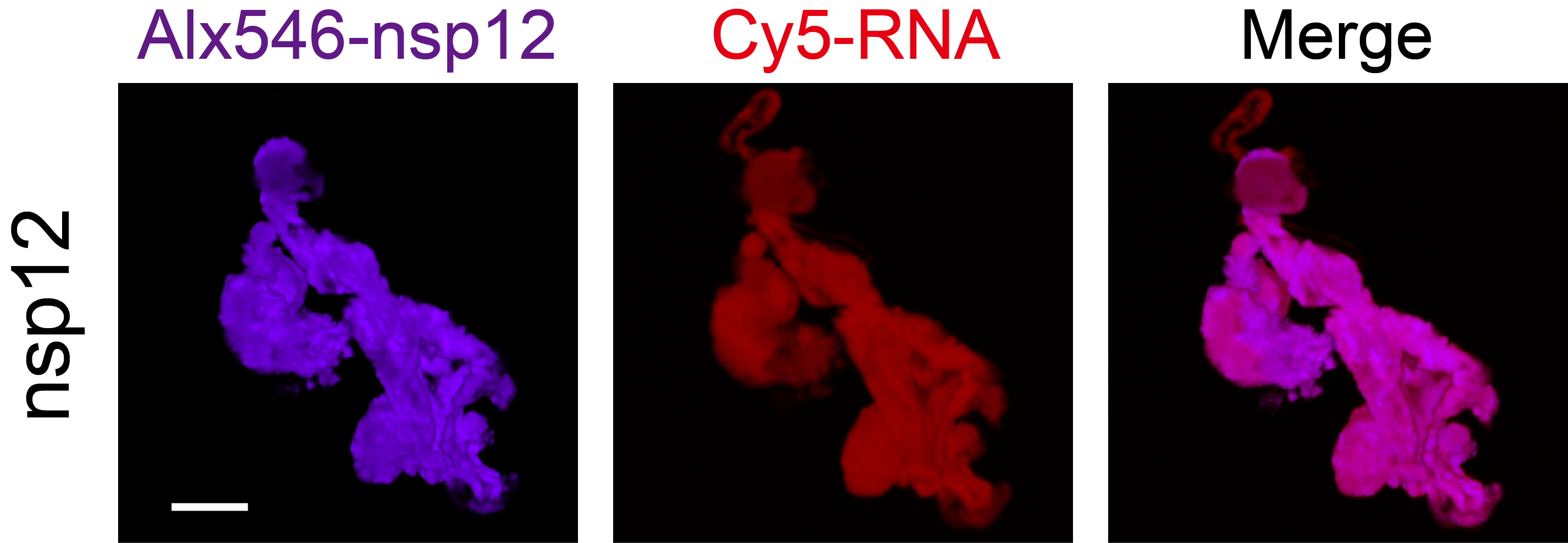

### Extended Data Fig5.png

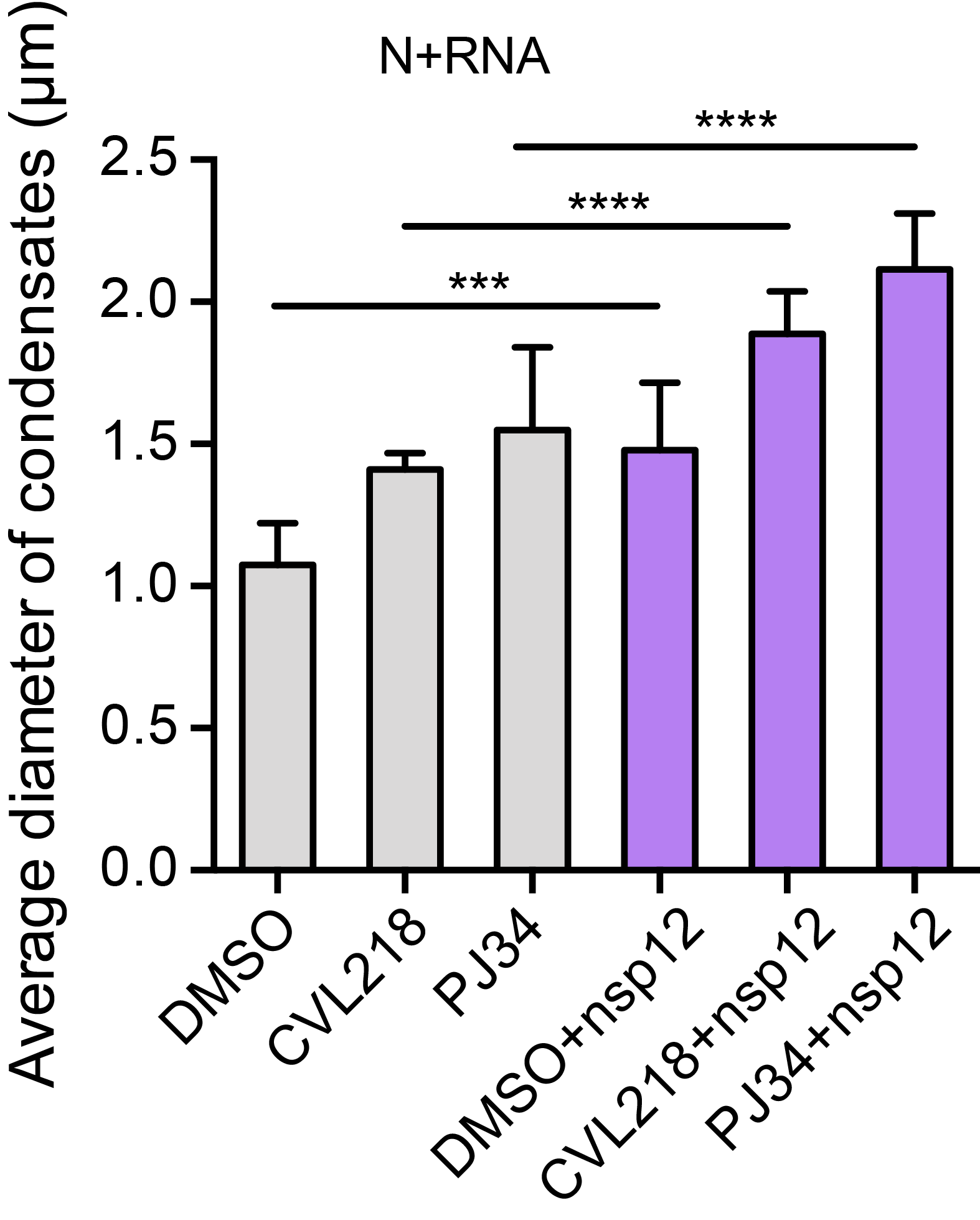

### Extended Data Fig6.png

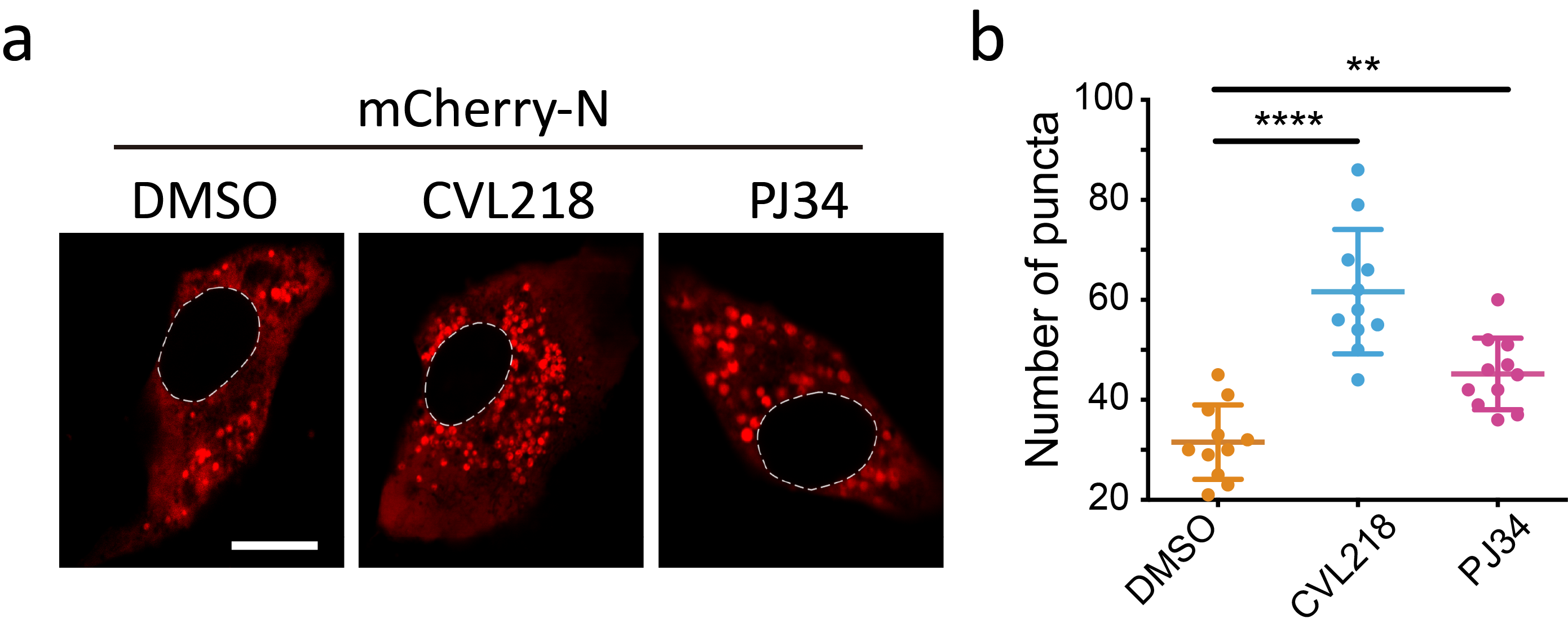
